# Genomic tailoring of autogenous poultry vaccines to reduce *Campylobacter* from farm to fork

**DOI:** 10.1101/2023.11.09.566360

**Authors:** Jessica K. Calland, Maiju E. Pesonen, Jai Mehat, Ben Pascoe, David J. Haydon, Jose Lourenco, Evangelos Mourkas, Matthew D. Hitchings, Roberto M. La Ragione, Philip Hammond, Timothy S. Wallis, Jukka Corander, Samuel K. Sheppard

**Affiliations:** Oslo Centre for Biostatistics and Epidemiology, Oslo University Hospital, Oslo, Norway; School of Biosciences, University of Surrey, Surrey, UK; Centre for Genomic Pathogen Surveillance, Big Data Institute, University of Oxford, Oxford, UK; Ineos Oxford Institute, Department of Biology, University of Oxford, Oxford, UK; Faculty of Veterinary Medicine, Chiang Mai University, Chiang Mai, Thailand; Ridgeway Biologicals Ltd. a Ceva Santé Animale Company, Berkshire, UK; Department of Zoology, University of Oxford, Oxford, UK; Swansea University Medical School, Swansea University, Swansea, U; School of Veterinary Medicine, University of Surrey, Surrey, UK; Crowshall Veterinary Services, Norfolk, UK; Oslo Centre for Biostatistics and Epidemiology, University of Oslo, Oslo, Norway; Department of Mathematics and Statistics, Helsinki Institute for Information Technology, University of Helsinki, Helsinki, Finland; Parasites and Microbes, Wellcome Sanger Institute, Cambridge, UK

**Keywords:** Autogenous vaccine design, Genomics, Campylobacter

## Abstract

*Campylobacter* is a leading cause of food-borne gastroenteritis worldwide, linked to the consumption of contaminated poultry meat. Targeting this pathogen at source, vaccines for poultry can provide short term caecal reductions in *Campylobacter* numbers in the chicken intestine. However, this approach is unlikely to reduce *Campylobacter* in the food chain or human incidence. This is likely as vaccines typically target only a subset of the high strain diversity circulating among chicken flocks and rapid evolution diminishes vaccine efficacy over time. To address this, we used a genomic approach to develop a whole-cell autogenous vaccine targeting isolates harbouring genes linked to survival outside of the host. We hyper-immunised a whole major UK breeder farm to passively target offspring colonisation using maternally-derived antibody. Monitoring progeny, broiler flocks revealed a near-complete shift in the post-vaccination *Campylobacter* population with a ∼50% reduction in isolates harbouring extra-intestinal survival genes and a significant reduction of *Campylobacter* cells surviving on the surface of meat. Based on these findings, we developed a logistic regression model that predicted that vaccine efficacy could be extended to target 46% of a population of clinically relevant strains. Immuno-manipulation of poultry microbiomes towards less harmful commensal isolates by competitive exclusion, has major potential for reducing pathogens in the food production chain.

## Introduction

Poultry meat has risen to become one of the main sources of affordable food protein worldwide^1^. Since the 1960s the global poultry flock has increased to approximately 26 billion birds^2^. While this has brought about nutritional benefits for humans, it has also come at a significant cost. Safe food production is among the most pressing challenges to sustainable agriculture and contamination of retail poultry is a critical issue. Retail poultry meat is the primary source of the common human enteric pathogens *Campylobacter jejuni* and *Campylobacter coli*^3–8^. These pathogens are typically introduced at slaughter^9,10^, with approximately 70% of retail poultry^11^ carrying these bacteria on their surfaces, often at concentrations exceeding 1,000 cells per gram of chicken skin. This is a concern when a single chicken breast can contain more than 30,000 *Campylobacter* cells, which is 60 times the minimum dose required for human infection^11,12^.

The most common advice for avoiding infection is careful handling and thorough cooking of poultry meat. However, it is questionable whether this is a sufficiently robust response to control a dangerous pathogen that enters kitchens and is responsible for acute gastroenteritis and debilitating and life threatening sequelae^13^. Numerous measures have been employed to reduce *Campylobacter* entering the food chain^14^. Broadly, these can be categorised as on-farm measures such as enhanced biosecurity^15^, bacteriocin treatments^16^, probiotics and feed supplements^17–22^, phage therapy^23^; or carcass treatments such as irradiation and chemical sterilisation^24^, increased hygiene to prevent intestinal contents spillage^15^, and hot water and freezing therapies. Although farm and factory-level preventative measures can reduce *Campylobacter* load, these have not yet translated to viable commercially relevant interventions in broiler production^15^.

Vaccination is an effective strategy to control or prevent infectious disease, and the immunisation of chickens with vaccines using inactivated bacteria have reduced intestinal *Campylobacter* colonisation by up to hundredfold (2 x Log10)^25^. However, despite multiple attempts^26–30^ an effective commercial vaccine to control *Campylobacter* in poultry has yet to be developed. The principal impediment is that *Campylobacter* populations in chickens can be very genetically and phenotypically diverse^31–36^ with multiple strains found together in a single flock, or even a single bird^34,37^. Therefore, inducing an immune response that will cross-protect birds against multiple strains is extremely difficult. As a result, even if a fraction of the population can be removed, vaccine escape strains can proliferate to fill the vacant niche space. One solution is to regularly update the vaccine to target prevalent strains, which is a common approach employed with the use of autogenous vaccines^38^. Widely applied by veterinarians, autogenous vaccines are typically prepared from pathogen isolates sampled on a particular farm, and then used to elicit strain-specific immune responses against homologous antigens in livestock on the same farm^38^.

The availability of increasingly large bacterial genome collections has greatly advanced understanding of variation in bacterial populations^39^, strain interactions^40^ and the function of specific genes in *Campylobacter* populations^41,42^. This has potential for improving vaccine development and delivery^43^. Specifically relating to *Campylobacter*, recent comparative genomic analyses of isolates from the intestines of broiler birds, carcasses and human clinical samples have shown that those infecting humans are a genetic subset of those found in poultry^41^. In particular, the genomes of the clinical isolates are enriched with genes that allow these microaerophilic organisms to tolerate atmospheric oxygen conditions and survive outside the gut^41,44^. Therefore, rather than trying to eradicate this ubiquitous commensal organism from the chicken gut, it may be possible to target only the strains that survive on food and in the environment, which ultimately infect humans.

Here we combine knowledge of *Campylobacter* strain variation in chickens and the genes that promote survival through poultry processing to inform autogenous vaccine design. Our passive immunisation approach aims to manipulate the gut microbiome to reduce clinically important *Campylobacter* and promote the proliferation of isolates that have less chance of surviving to contaminate retail meat^45^. Hyper-immunising chickens at a major UK breeder farm, we target a relatively brief *Campylobacter* colonisation window, 14 days after hatching when maternally-derived immunoglobulin (Ig) Y antibodies that protect young birds^46^ begin to dwindle, but before slaughter at around 37 days. Tracking chickens through slaughter, we investigate the potential for genomically-informed autogenous vaccines as a potential near real-time treatment to reduce *Campylobacter* in poultry, and ultimately human infections.

## Results

### Broiler chickens sampled pre-vaccination harbour multiple Campylobacter lineages

A total of 150 caeca and 150 neck skin samples were collected from 300 Ross broiler chickens across five broiler farms in the UK. From these, *C. jejuni* was cultured from 189 (101 neck skin, 88 caeca) and 136 whole-genomes (74 neck skin, 62 caeca) were sequenced (S1 and S2 Table). Phylogenetic comparison with other chicken-associated lineages revealed four distinct clonal complexes (CCs) (Fig 1A): CC257 (n = 53), CC443 (n = 26), CC206 (n = 44) and sequence type (ST) ST-4430 (n = 13), defined by PubMLST^47^ belonging to an undefined/mixed cluster (Fig 1A, S1 Table). Three of the four identified CCs (CC257, CC443 and CC206) were reported in human clinical cases in Oxford over a four-year period (S3 Table), consistent with studies indicating transmission from poultry to humans^5^.

**Fig 1.**
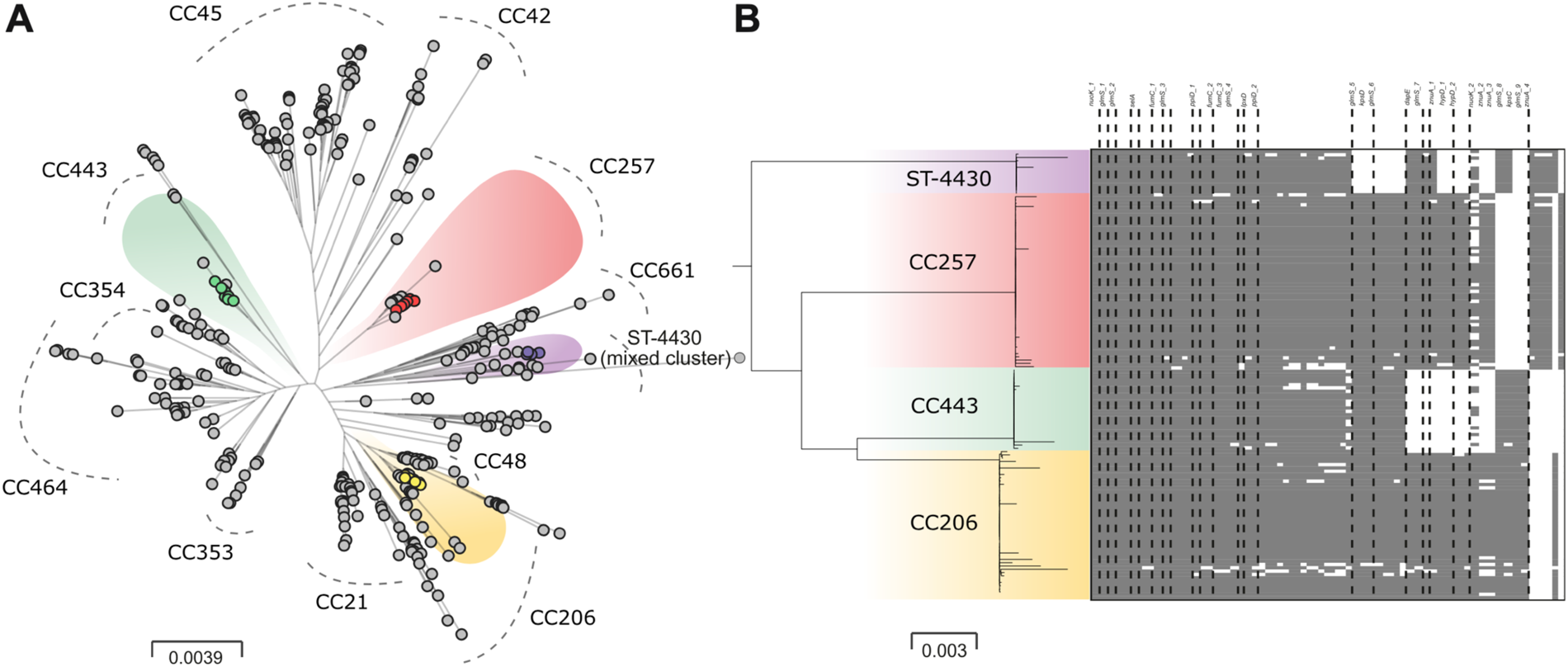
Highly structured *Campylobacter* populations inform vaccine design. (A) Maximum Likelihood (ML) phylogeny of 1,095 *C. jejuni* genomes indicating major chicken-associated CCs (grey dashed curves) and isolates in the pre-vaccination population (n = 136, faded halos). Scale bar represents substitutions per site. (B) Presence/absence (grey/white cells) distribution of 70 survival-associated genetic elements in pre-vaccination genomes CCs ordered by the frequency within all genomes. Only genetic elements with an annotated gene name are labelled (complete list in S4 Table). Variation of a genetic element with the same gene annotation are labelled as ‘_1’, ‘_2’, ‘_3’ etc. The matrix rows are ordered by the genome position in the phylogeny left of the presence-absence matrix.

### Elements linked to environmental survival were identified in C. jejuni isolates

The frequency of genes that promote the survival of micro-aerophilic *Campylobacter* outside of the host provide information about the likelihood that a given isolate will persist through the poultry production chain^41^. Known survival-associated genetic elements were identified in *C. jejuni* genomes from unvaccinated chickens (S4 Table, Fig 1B). The average number varied by CC as follows: CC257 (n = 65), CC443 (n = 60), CC206 (n = 66), ST-4430 (n = 61). The number of survival-associated elements also varied within CCs: CC257 (51-67), CC443 (52-63), CC206 (47-68), ST-4430 (46-63). Four isolates (one from each CC) containing the most survival-associated genetic elements were chosen for inclusion in the vaccine (isolate ids: 7939276 (ST-4430), 7939216 (CC257), 7916660 (CC443), 7930870 (CC206)) based on the likelihood of these isolates surviving outside the host gut.

### Campylobacter-vaccinated birds developed a strong strain-specific IgY response

Immunoglobulin Y is the major immunoglobulin (Ig) class in chickens. This is passed to the embryo from the breeder hen via the egg yolk and protects chicks against *Campylobacter* colonisation^48,49^. Strain-specific IgY titres were quantified in breeder and progeny broiler serum and breeder egg yolk (Fig 2). In breeder serum, the specific IgY response to the four-strain combination vaccine was significantly higher (*P* = ≤0.05) compared with the control breeders at all three time points (TPs). Specifically, mean OD_450_ of 8.026 (range 1.651 – 2.513) vs 0.535 (0.131 – 0.137), 15.658 (3.823 – 3.950) vs 0.515 (0.122 – 0.133), 13.720 (3.426 – 3.437) vs 0.522 (0.128 – 0.136) for time points 1, 2 and 3 respectively (Fig 3A, S1 Fig). IgY remained consistently low in all control cohorts (∼0.500). There was a reduction in total vaccine-specific IgY titres from vaccinated breeder hens in the breeder egg yolk (mean OD_450_ = 2.206) vs breeder blood serum (mean OD_450_ = 12.468) but increased slightly in control cohorts (breeder serum mean OD_450_ = 0.524, breeder egg yolk mean OD_450_ = 1.131). Vaccine-specific IgY maternally-derived antibody was highest at TP1 in the vaccinated breeder egg yolk samples vs the control group with mean OD_450_ of 4.362 (0.310 – 2.456) vs 0.510 (0.083 – 0.171). Mean OD_450_ reduced by TP2 in vaccinated (0.689 (0.034 – 0.247)) vs control groups (1.310 (0.234 – 0.415)) and increased slightly by TP3 (1.568 (0.132 – 0.532) vs 1.573 (0.336 – 0.434), for vaccinated and control cohorts respectively). In progeny broiler blood serum the mean total vaccine-specific IgY titres were higher than in egg yolk in both vaccinated (mean OD_450_ = 8.053) and control (mean OD_450_ = 8.675) cohorts and was present at all TPs at a mean OD_450_ of 9.132 (2.206 – 2.411) vs 12.210 (2.579 – 3.524), 4.634 (0.616 – 1.791) vs 5.793 (1.306 – 1.666) and 10.392 (2.357 – 3.052) vs 8.023 (1.794 – 2.225) (S1 Fig) for vaccinated vs control cohorts at TP1, TP2 and TP3 respectively. Antibody titres from both vaccinated and control broiler blood samples were highest at TP1 and reduced at TP2, consistent with the natural waning of maternally-derived antibodies (MDAs) (Fig 3C, S1 Fig).

**Fig 2.**
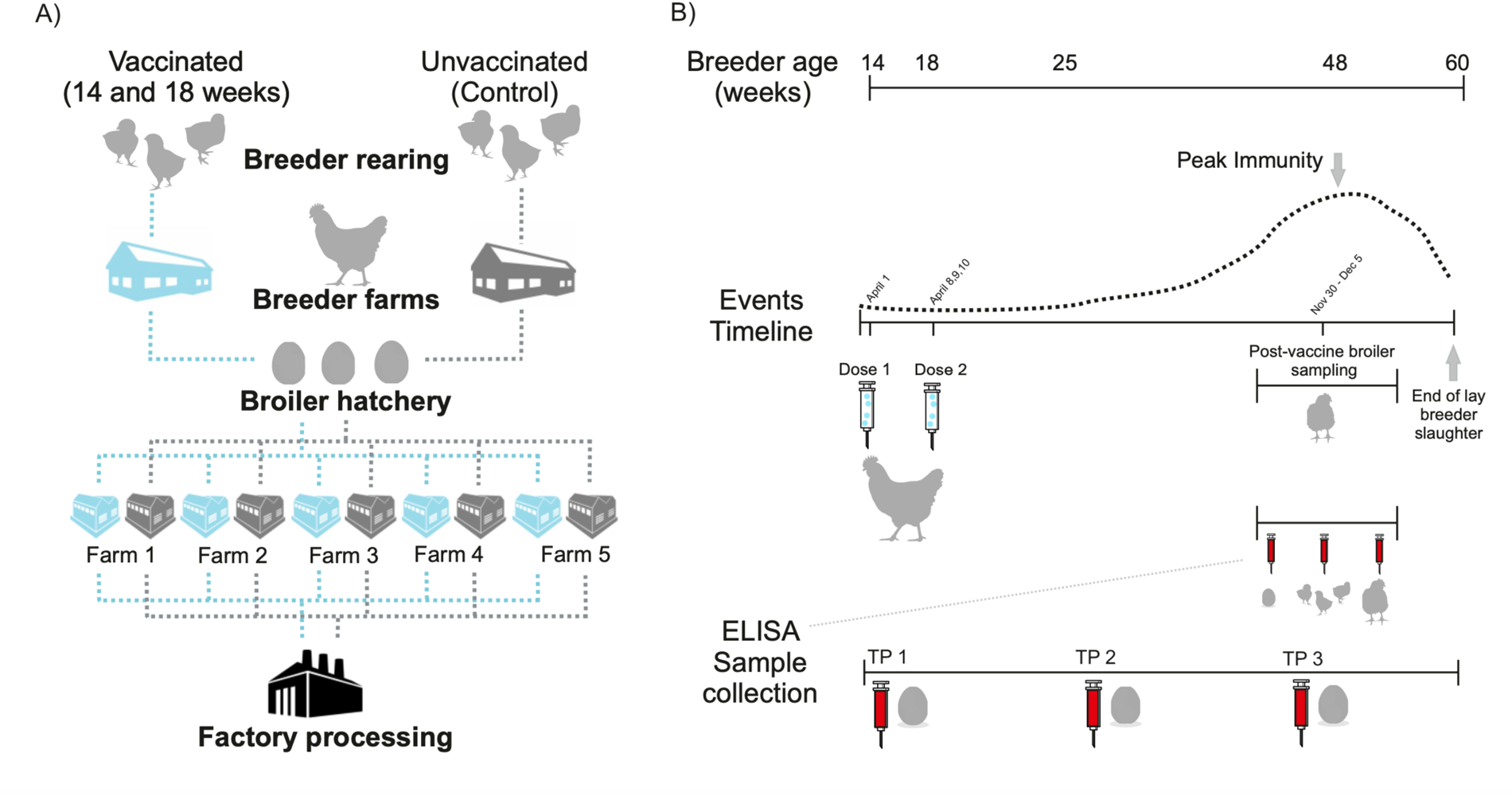
Cohort-controlled poultry distribution system and vaccine implementation. (A) Breeder birds go to farms at age of lay and progeny of are distributed to hatcheries as eggs and hatched broiler chicks are reared in broiler farms. Broilers remain in the same farm house until age of slaughter (∼37 days) and meat processing. Vaccinated breeders and progeny were followed as one epidemiological unit through a controlled system to ensure vaccinated (blue) and control (grey) cohorts remained separate (origin, hatchery, broiler houses, abattoir). (B) A whole breeder rearing farm (∼40,000 breeder birds) were immunised with two doses of the vaccine (14-18 weeks of age). Breeder eggs and blood were sampled (for ELISAs, IgY MAb) at three time points and *Campylobacter* were sampled at peak immunity age from progeny (vaccinated and control) caeca and neck skin.

**Fig 3.**
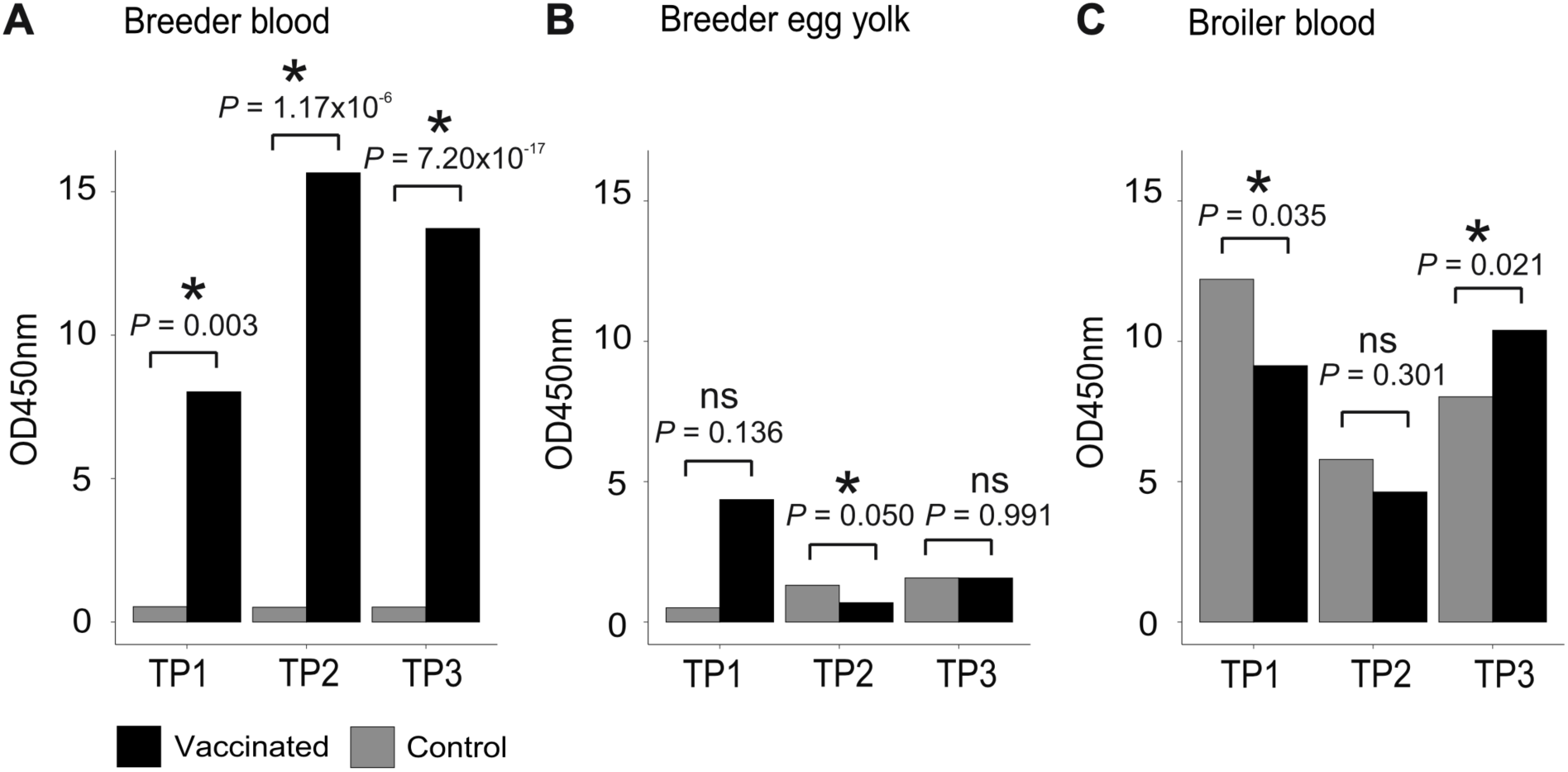
Vaccine-specific immunogenic effects in vaccinated birds. Bar charts of the average concentrations (at optic density, OD_450_) of vaccine-specific IgY from the four vaccine isolates, in vaccinated (black) and control (grey) breeder bird blood (A), breeder egg yolk (B) and broiler blood (C). Vaccine-specific IgY titres were measured for three time points (TP) across the breeder/broilers’ life. Statistical significance between vaccinated and control cohorts was measured using a two-sample t-Test and significant difference (P < 0.05) is indicated with an asterisk and non-significance by ‘ns’.

### Target Campylobacter isolates are replaced in the post-vaccination chicken gut

*Campylobacter* were grown from 667 samples (neck skin control (n = 97); neck skin vaccinated (n = 200); caeca control (n = 132); caeca vaccinated (n = 238), of which 117 were sequenced (S1 Table, Table 1, Fig 4). Phylogenetic comparison revealed an almost complete post-vaccination strain replacement with only 3 isolates sharing CCs with the pre-vaccination population. The remaining isolates (114 out of 117) sampled from the post-vaccination population belonged to three *C. jejuni* chicken-associated lineages: CC353 (n = 33), host generalist CC21 (n = 25), and ST-7735 (n = 4), a lineage with a high degree of shared ancestry with ST-4430 (sampled pre-vaccination) (Fig 4) and belonging to the same undefined/mixed CC. The fourth lineage sampled from the post-vaccination population was the *C. coli* host generalist CC828 (n = 52). The same CCs were present in both vaccinated and control flocks but at different frequencies from different sample types (Table 1). The post-vaccination shift in *Campylobacter* population structure is consistent with isolate replacement promoted by the vaccine (S1 Table, Table 1, Fig 4A).

**Fig 4.**
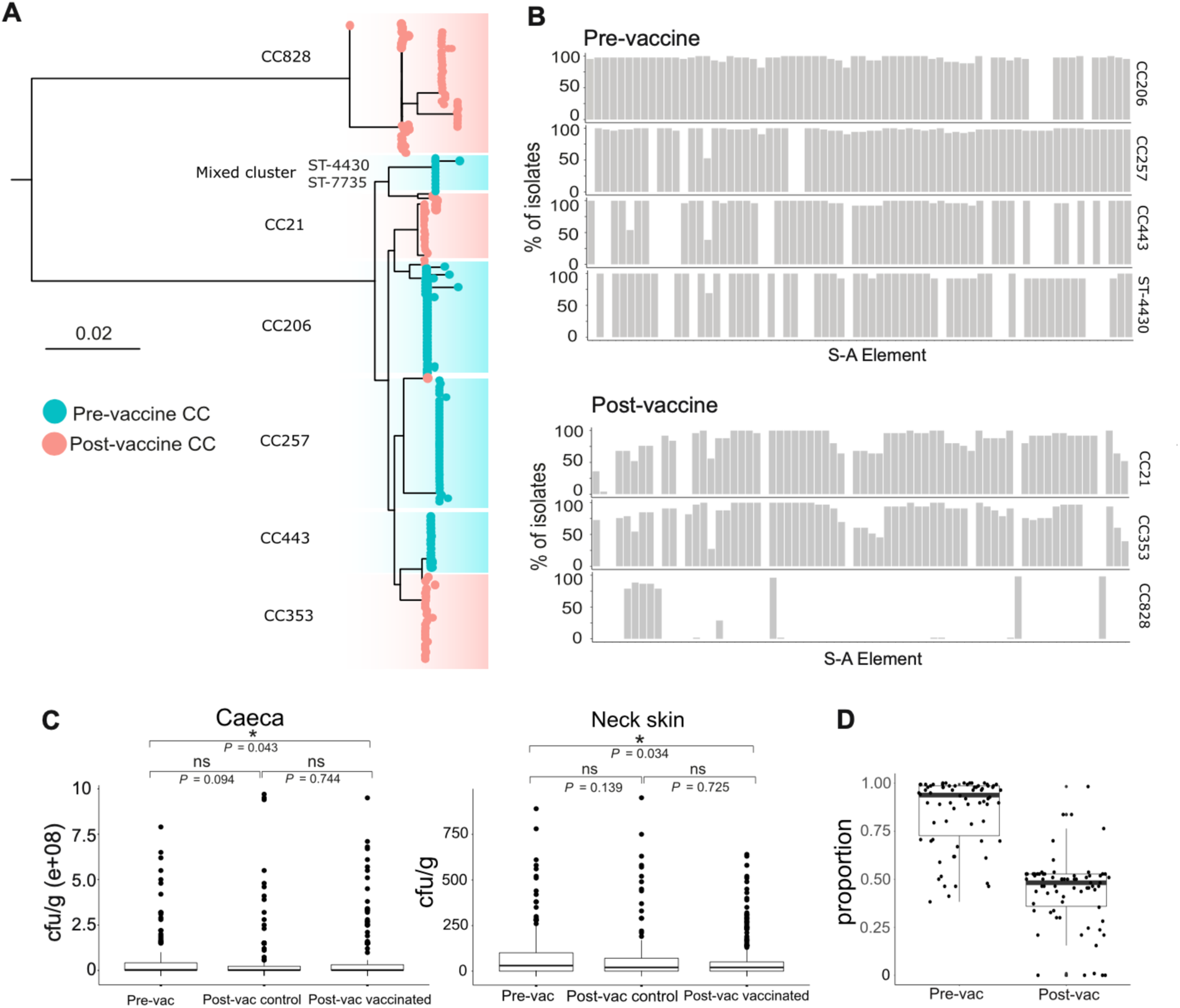
Targeted passive vaccination reduces *Campylobacter* enriched with survival-associated genes. (A) Phylogenetic comparison of pre-(blue) and post-(pink) vaccination genomes from each clonal complex (CC). The scale bar represents substitutions per site. (B) Histograms indicate the % of isolates containing specific survival-associated (S-A) elements (S4 Table) within clonal complexes (CCs) found in pre-(above) and post-(below) vaccine populations. ST-7735 is not shown due to the low sample size (n=4). (C) Boxplots of *Campylobacter* levels (cfu/g) in caeca and on neck skin samples are shown for pre- and post-vaccine and control groups and pairwise comparisons of means *p-*values (Mann-Whitney U tests, P <0.05) are shown with significance indicated by an asterisk and non-significance by ‘ns’. Overall significance was estimated using the Kruskal-Wallis test. (D) The overall proportion of isolates containing a survival-associated elements (black dots) are lower in the post-vaccination population.

**Table 1.** Comparison of CCs between pre- and post-vaccination genomes.

| CCs | Pre-vac |  | Post-vac |  |  |  |  |
| --- | --- | --- | --- | --- | --- | --- | --- |
|  | neck skin | ceaca | vaccinated |  | control |  | source unknown |
|  |  |  | neck skin | ceaca | neck skin | ceaca |  |
| CC206 | 21 | 23 | - | - | - | - | - |
| CC257 | 31 | 22 | - | 1 | - | - | - |
| CC443 | 10 | 16 | 2 | - | - | - | - |
| ST-4430* | 12 | 1 | - | - | - | - | - |
| ST-7735* | - | - | - | - | 1 | 2 | 1 |
| CC21 | - | - | 17 | 3 | - | 1 | 4 |
| CC353 | - | - | 7 | 19 | 1 | 6 | - |
| CC828 | - | - | 12 | 13 | 14 | 12 | 1 |

\*belonging to closely related (mixed) cluster

### There are fewer survival-associated genes in post-vaccination Campylobacter genomes

The frequency of specific survival-associated genetic elements differed between CCs and between pre- and post-vaccination populations (Fig 4B and D). The range in the number of survival-associated genetic elements per isolate varied by CC (ST-4430 = 38-52, CC206 = 45-65, CC257 = 49-65, CC443 = 42-53, CC21 = 29-61, CC353 = 17-59, CC828 = 1-14, ST-7735 = 28-55) (Fig 4B). This corresponds to a combined average of 226 per pre-vaccination genome, reducing to 110 in the post-vaccination population. The reduction is even more striking when comparing the total number of survival-associated elements, with a total of 8,098 total survival elements in the pre-vaccination population versus 3,377 in post-vaccination isolates (Fig 4D). The presence of survival-associated elements differed among CCs. For example, only 6 out of the 70 survival-associated elements were present in all CCs (13_*Cj1049c*, 14_*Cj1049c*, 15_*Cj1049c*, 16_*Cj1049c*, 5_IpxD_*Cj0576*, 6_*hypD*_*Cj0625*) but at varying proportions (Fig 4B). This means that 64 of the 70 element types were not shared by all of the CCs in both populations. However, 41 of the survival-associated elements were present in all isolates in the pre-vaccination population and 10 in all isolates in the post-vaccination population (Fig 4B).

### Vaccination does not reduce caecal load but lowers numbers of Campylobacter surviving on meat

*Campylobacter* were enumerated in cfu/g for each caeca (n = 88) and neck skin (n= 101) sample pre-vaccination (Fig 4C and Table 2). Levels of positive *Campylobacter* growth in the caecum varied across the five farms sampled and ranged from 7-100%, with one farm testing completely negative for *Campylobacter* in the caeca (Table 2, S6 and S7 Tables). Any caecal sample containing <1000 cfu/g was classed as *Campylobacter* negative (S2 Table) according to standard laboratory protocol (Jorgensen et al., 2018). Caecal load for *Campylobacter* ranged from 2.95E+06 – 2.05E+08 cfu/g, with an average of 6.1E+07 cfu/g (Fig 4C, S2 Fig and Table 2). Enumeration of *Campylobacter* on neck skin samples also differed per farm (Farm 1 = <10-4500; Farm 2 = <10-70; Farm 3 = <10-3700; Farm 4 = <10-80; Farm 5 = <10-32000 cfu/g) (S6 and S7 Table). However, for the neck skin samples, anything <10 cfu/g were classed as negative according to standard laboratory protocol and anything >1000 cfu/g were classed as outliers and removed (S6 and S7 Tables, S2 Fig). Average *Campylobacter* enumeration on the neck skin was estimated at 150.0 cfu/g across all farms once exclusion criteria had been applied (Farm 1 = 0-610; Farm 2 = 0-32; Farm 3 = 0-147; Farm 4 = 0-80, Farm 5 = 0-780 cfu/g) (Table 2, S6 and S7 Tables, Fig 4C).

**Table 2.**
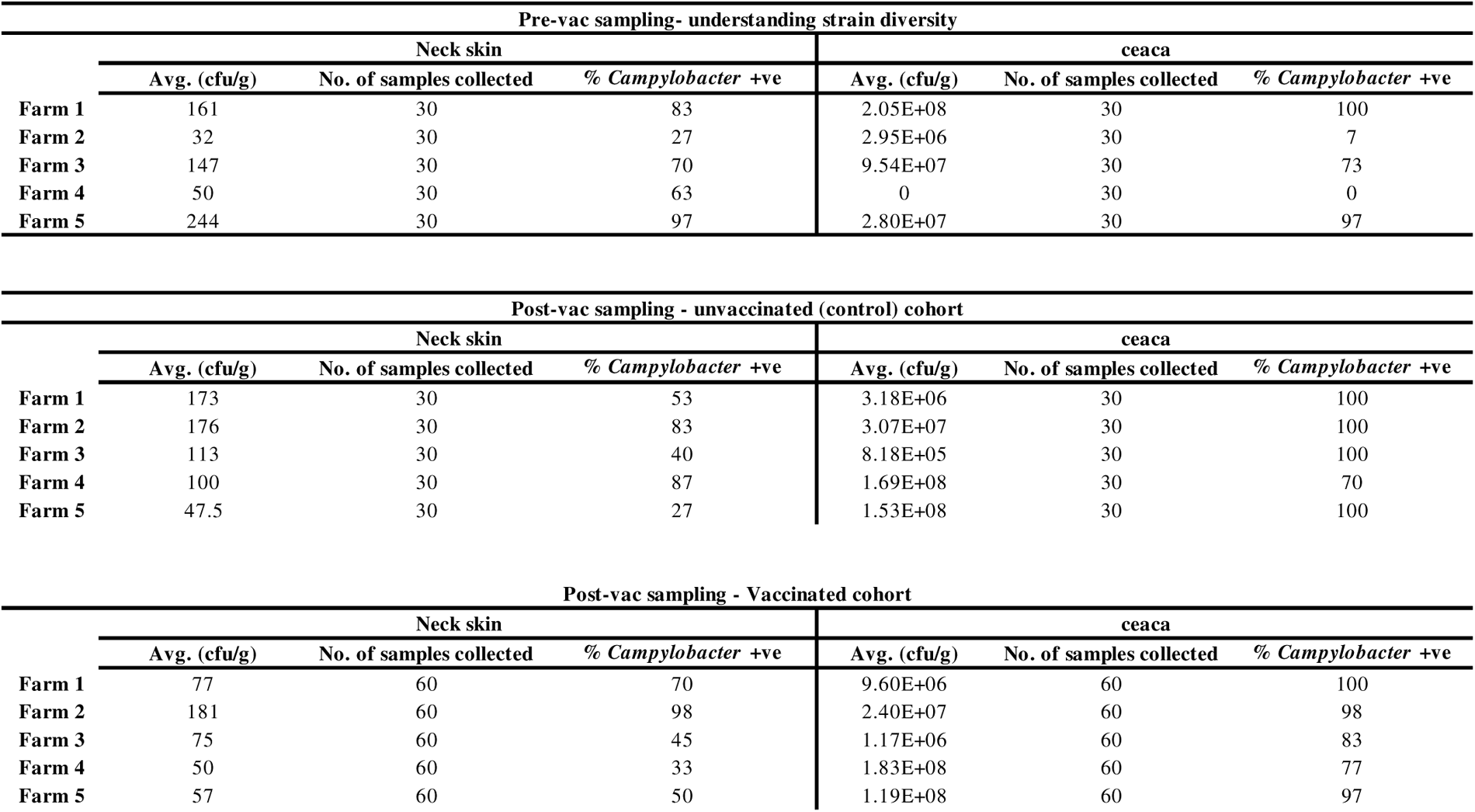
Per-farm *Campylobacter* prevalence summary pre- and post-vaccination.

| Pre-vac sampling- understanding strain diversity |  |  |  |  |  |  |
| --- | --- | --- | --- | --- | --- | --- |
|  | Neck skin |  |  | ceaca |  |  |
|  | Avg. (cfu/g) | No. of samples collected | % *Campylobacter* +ve | Avg. (cfu/g) | No. of samples collected | % *Campylobacter* +ve |
| Farm 1 | 161 | 30 | 83 | 2.05E+08 | 30 | 100 |
| Farm 2 | 32 | 30 | 27 | 2.95E+06 | 30 | 7 |
| Farm 3 | 147 | 30 | 70 | 9.54E+07 | 30 | 73 |
| Farm 4 | 50 | 30 | 63 | 0 | 30 | 0 |
| Farm 5 | 244 | 30 | 97 | 2.80E+07 | 30 | 97 |

| Post-vac sampling - Vaccinated cohort |  |  |  |  |  |  |
| --- | --- | --- | --- | --- | --- | --- |
|  | Neck skin |  |  | ceaca |  |  |
|  | Avg. (cfu/g) | No. of samples collected | % *Campylobacter* +ve | Avg. (cfu/g) | No. of samples collected | % *Campylobacter* +ve |
| Farm 1 | 77 | 60 | 70 | 9.60E+06 | 60 | 100 |
| Farm 2 | 181 | 60 | 98 | 2.40E+07 | 60 | 98 |
| Farm 3 | 75 | 60 | 45 | 1.17E+06 | 60 | 83 |
| Farm 4 | 50 | 60 | 33 | 1.83E+08 | 60 | 77 |
| Farm 5 | 57 | 60 | 50 | 1.19E+08 | 60 | 97 |

Enumeration of *Campylobacter* isolates present in caeca samples across five farms post-vaccination showed little difference in average cfu/g between the control (6.0E+07), and vaccinated (5.6E+07) cohorts (non-parametric Mann-Whitney U-tests, *P* = 0.744 (not significant)). However, a significant difference between pre-vaccination (6.1E+07) and post-vaccination vaccinated cohorts was observed (*P* = 0.043) (Fig 4C and Table 2). After the removal of outliers as in pre-vaccination populations (>1000 cfu/g) (S6 and S7 Tables, S3 Fig), there was a reduction in average neck skin counts from 150.0 cfu/g (pre-vaccination) to 118.0 cfu/g in the control cohorts (post-vaccination). The reduction of surviving *Campylobacter* was greater on vaccinated neck skin samples (92.0 cfu/g) (Fig 4C and Table 2). Pairwise comparison of all groups showed a significant reduction between the pre-vaccination and post-vaccination neck skin cohorts (non-parametric Mann-Whitney U-tests, *P* = 0.034).

### Genome comparison can be used to model vaccine efficacy in complex Campylobacter populations

As a whole-cell vaccine was developed, there are limitations identifying the specific antigens underlying the targeted immunogenic response and if this immunogenic effect will confer cross-protection to the antigens of other lineages or CCs. We implemented a logistic regression model-based approach using antigenic variation from the genomes from this study to better describe this and to predict the effects of our vaccine isolates at targeting other chicken-associated CCs in a hypothetical population. First, the average fraction of the relative frequencies of the vaccine isolates present post-vaccination was computed, resulting in a baseline probability (intercept of the model) of 0.08. In simple terms, this can be interpreted as an 8% chance of an isolate identical to a vaccine isolate (at the selected antigenic loci) surviving the effects of the vaccine and ending up in the post-vaccination population. Next, three different scenarios were created to assess the effect of the vaccine: logistic regression model parameter thetas were fixed to 1.099, 0.916 and 0.693, which correspond to odds ratios (OR) 3, 2.5 and 2, respectively (Fig 5B, S4 Fig). An OR of 2.5 was selected for further analysis. This provides an indication of how distinct antigens could be in order for the isolate to escape the effects of the vaccine as a function of the distance. For example, according to the model, antigen sequences can differ by ∼2000 amino acids (8% of antigen sequence divergence) from the vaccine isolates to increase likelihood of survival by ∼40% and by ∼8000 amino acids to result in a predicted ∼100% likelihood of surviving the effects of the vaccine (Fig 5B, S4 Fig) (31% of vaccine antigen sequence divergence). These factors combined suggest evidence for high likelihood of strain replacement.

**Fig 5.**
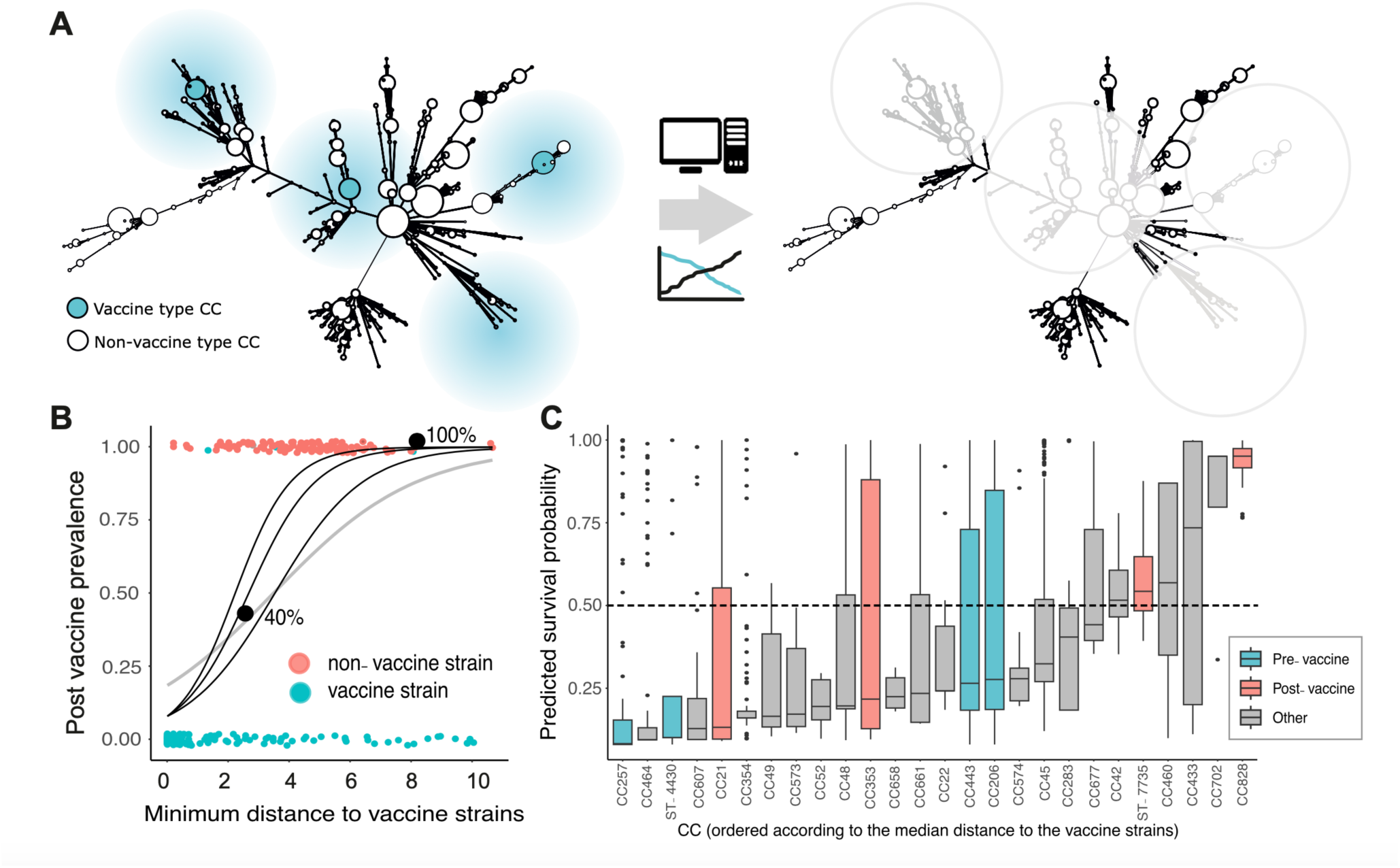
Inferring vaccine efficacy in complex *Campylobacter* populations. (A) Grape diagrams showing the predicted zone of efficacy for vaccine isolates (blue), inferred for common *Campylobacter* clonal complexes (CCs) based upon allelic similarity and the predicted post-vaccine population is shown (right). (B) The logistic regression model (grey curve) fitted to the trial data to predict the probability of isolates surviving the effects of the vaccine isolates (y axis). Black curves represent different thetas (1 = 1.099 (OR = 3); 2 = 0.916 (OR = 2.5); 3 = 0.693 (OR = 2)) and coloured circles indicate the minimum distance of each isolate to one of the four vaccine isolates (x axis). Blue and pink circles belong to the CCs found predominantly in pre-(vaccine isolate CCs) post-vaccination populations. Likelihood of escaping the vaccine is indicated by back circles. (C) Strain replacement potential was estimated from the distribution of isolate survivability (black dots) probabilities per chicken-associated CC – ordered by median survivability (black horizontal lines). There was variation in inferred isolate survivability within CCs, including pre-(pink) and post-(blue) vaccine sampling (pink), but they can be ordered from least to most likely to escape vaccine effects and comparison to dashed horizontal line (0.5) indicates the minimum probability at which survival post-vaccine is likely.

The model results were then used to predict the effects of the vaccine isolates in a wider context using a hypothetical population of chicken-associated isolates. A survivability probability was computed for each minimum distance of the context chicken isolate collection to the vaccine isolates using the results from the model (theta = 0.916, OR 2.5). The isolates were then grouped by CC to result in 31 identified CCs (S3 table). Any CC containing <3 sequences were removed (CC362, CC581, CC446, CC61 and CC508), leaving 26 CCs for analysis. The distributions of probability of vaccine survival for each isolate in each CC were assessed (Fig 5). Considerable variation was observed for predicted survivability between CCs as a whole but a positive trend for escaping the effects of the vaccine with increased distance from the vaccine antigens was clear. Of the 26 CCs analysed, only 6 exhibited a median predicted survival probability >0.5 (CC42 = 0.516, ST-7735 = 0.543, CC460 = 0.569, CC433 = 0.735, CC702 = 0.951 and CC828 = 0.951) (>0.5 suggests a high likelihood of escaping the effects of the vaccine). Two of which were present in the post-vaccination population (replacement strains) (ST-7735 and CC828). *C. coli* CC828 had the highest survival probability score of all the CCs (Fig 5).

Of the remaining CCs analysed, 8 exhibited a range in survival which spread >0.5 (CC677, CC45, CC206, CC443, CC661, CC353, CC48 and CC21) suggesting high likelihood and variability of survival within the CC. Two of these CCs (CC21 and CC353) belonged to the remaining CCs sampled post-vaccination in the vaccine trial (Fig 5) and two belonged to CCs which were included in the vaccine (CC443 and CC206). The remaining 12 CCs (CC257, CC464, ST-4430, CC607, CC354, CC49, CC573, CC52, CC658, CC22, CC574, CC283) had both median predicted survival probabilities <0.5 and narrow ranges in survivability for the CC as a whole, suggesting that these CCs would likely be targeted by the vaccine and not survive post-vaccination. Two of these (CC257 and ST-4430) were CCs included in the vaccine. In this case, we can speculate that in a hypothetical population with 26 distinct circulating *Campylobacter* CCs, our vaccine isolates may be effective at targeting (knocking out) 12 out of 26 CCs which corresponds to 46% of the population.

## Discussion

Vaccines are highly effective in reducing both infections and the need to use antibiotics for treatment of bacterial infections, which explains their popularity in both human and agricultural practices. However, there are significant challenges associated with effective design for targeting genetically and antigenically diverse organisms, such as *Campylobacter* in chickens. Developing a commercially viable on-farm poultry vaccine has proven elusive, despite the clear benefits of combating strains that contaminate meat and cause human campylobacteriosis. Various vaccine types have been previously trialled to reduce *Campylobacter* load in poultry, such as conjugate, subunit, whole-cell, *in ovo*, and egg yolk IgY routes^29,30,46,50–54^. However, the poorly understood chicken cellular response to this ostensibly commensal bacterium complicates traditional vaccine approaches^55^. This led us to prioritise population manipulation by delaying colonisation of isolates likely to survive on meat, rather than relying solely on vaccine-induced cellular responses. Using genomic approaches and rational design targeting isolates surviving processing, we designed and manufactured a farm-specific autogenous vaccine and trialled its efficacy in real time through a poultry distribution and processing system in the UK and monitored the effects over two years.

Leveraging the window in the first two weeks of a broiler’s life, where *Campylobacter* levels are low due to the protective maternal antibodies (IgY) passed from mother to offspring^17,46,48,49,56^, our passive vaccine approach facilitates vaccine administration, with whole cell *C. jejuni* vaccination in breeder hens conferring the protective effects of maternal antibodies (IgY) to offspring via the egg yolk to immunise chicks^50,52,54^. ELISA’s showed that vaccine-specific IgY was induced in vaccinated breeders and was passed to offspring through eggs despite peaking earlier than in a comparable study^57^ (Fig 3A, S1 Fig). As expected, protective maternal antibody levels waned after two weeks in the broiler serum (Fig 3C) coinciding with the natural *Campylobacter* colonisation^46^, but vaccine-specific IgY remained elevated in both vaccinated and control cohorts. This was potentially due to cross-reactivity with naturally acquired lineages from the farm^58,59^. Genomic characterisation of post-vaccination isolates revealed the small presence of pre-vaccination isolates circulating on the farms (Table 1) and could offer further explanation to the presence of vaccine-specific IgY in control cohorts^59^.

Consistent with the population manipulation approach, live-attenuated oral vaccines can also be tailored to target high-risk aerotolerant isolates as in mutagenesis studies^60^. Vaccine design was informed by prior knowledge of the genes linked to survival through poultry production^41^, and included a single isolate from each of the four pre-vaccination lineages with the highest predicted survivability. By selecting for isolates predicted to be less likely to survive when shed into the environment, we aimed to reduce persistence through transit to infect humans or contaminated poultry meat. As expected, these lineages were replaced in vaccinated chickens and, importantly, the post-vaccine population had on average fewer survival elements. While the caecal isolate composition changed, the overall caecal load was not significantly different in vaccinated and unvaccinated chickens. However, there was a substantial reduction in *Campylobacter* on neck skin at slaughter in vaccinated poultry cohorts, falling as low as 92 cfu/g.

The replacement of vaccine-type strains with non-vaccine types (NVTs) at the population level is a serious problem for infection control and has been observed among asymptomatic carriers of *Streptococcus pneumoniae* following the widespread use of the (PCV7) conjugate vaccine designed to target seven of the >92 serotypes^61,62^. Our empirical logistic regression model assessed the likelihood of the observed strain replacement, focusing on antigenic variation between pre- and post-vaccination CCs. Notably, CC828, which is the dominant *C. coli* clonal complex had the highest median survival probability with other replacement CCs (ST-7735, CC353 and CC21) exhibiting a high predicted likelihood of replacement. This is not surprising as the vaccine consisted of only *C. jejuni* isolates but highlights the potential importance of *C. coli* isolate inclusion in future vaccine design. When applied to a comprehensive collection of known chicken-associated CCs, the model revealed those that would likely survive in post-vaccination populations. Given the strain-specific nature of immunogenic responses and the unique antigen content of individual sequence types within CCs, differences were anticipated. However, modelling potential vaccine impact based on antigenic similarity in diverse chicken *Campylobacter* populations predicted that 12 out of 26 CCs would be targeted by the vaccine, representing 46% of the population. Hence there is an opportunity for future modelling studies to investigate optimisation of the vaccine composition based on genomic surveillance of the local *Campylobacter* population.

However, there were limitations to our study design. Four isolates containing a maximal number of survival-associated genetic elements from each of the four lineages identified from the pre-vaccination sampling were selected to be included in the vaccine. The rationale being that these four isolates had the highest expected potential of survivability through the poultry processing chain among all isolates from the farm surveillance. The individual contribution of a single genetic element to the differential survival of a bacterial cell on poultry meat is unknown, and most likely dependent on the niche conditions of a particular stage during processing. Therefore, our choice of isolates may not be optimal, but opens up new research avenues involving functional characterisation and provides a basis for modelling vaccine impact. By including as many survival-associated elements as possible, it is anticipated that the likelihood of survival is increased by essentially preparing for as many environmentally stressful scenarios as possible.

In conclusion, our combined vaccine design and predictive modelling approach demonstrates the potential for incorporating population genomic surveillance into vaccine design strategies for controlling opportunistic pathogens with extensive genetic variation. Modelling approaches can make informed predictions of efficacy and offer potential for optimising vaccine composition. Finally, rather than seeking eradication, a vaccination strategy that specifically targets lineages associated with an elevated risk of transmission, persistence, and ultimately infections is a promising approach to combat pathogenic isolates that live within larger commensal populations.

## Materials and Methods

### Ethics

No chickens were infected with *Campylobacter*, blood-sampled or euthanised for the sole purpose of this study. At *post-mortem* examination, removal of the caeca and neck flap skin was carried out by a trained veterinarian who had obtained permission from farm owners. Sampling was discussed appropriately between researchers, veterinarians, industrial collaborators and farmers prior to sampling procedures. Vaccinations with the autogenous vaccine were carried out by trained personnel and overseen by a veterinarian.

### Isolate collection

Sampling was conducted in two stages, pre-vaccination and post-vaccination. The progeny from one vaccinated and one control (non-vaccinated) breeder farm were followed through production. For pre-vaccination sampling, *Campylobacter* isolates were sampled to estimate isolate variation in caeca and on neck skin. From these isolates, candidate “survivor” isolates were identified for inclusion in the vaccine. *Campylobacter* is found at highest concentrations in the caecum. Therefore, 30 caeca samples from 30x 34-38 day-old Ross broilers from a UK poultry company were sampled across two days from 5 broiler farms (150 samples) leading into one abattoir. Additionally, 30 neck skin samples were taken from birds from the same flocks after 6 days of refrigerated storage at 3°C (150 neck skin flaps). This replicated the conditions of packaged meat purchased in supermarkets and maximised the likelihood of identifying isolates that can survive on food products (S1 Table).

Post-vaccination sampling was conducted to determine the effects of the vaccine on *Campylobacter* isolate variation, both within the broiler gut and on isolates persisting on poultry meat. Vaccinated and control (non-vaccinated) populations were followed from the rearing stages in separate cohorts (Fig 2). Sixty caeca from 60 x 37-day old broiler birds from each of the 5 broiler farms were sampled from the vaccinated progeny at peak breeder immunity age (∼48 weeks of age). This was approximately 30 weeks after the second dose of vaccine was administered (300 samples). In parallel, 60 neck skins were sampled from the same flocks 6 days after slaughter (300 samples). Thirty caeca samples were collected from each of the control population farms (150 samples) along with 30 neck skins from the same flocks after 6 days of refrigerated storage at 3°C (150 samples).

### Culture and isolation

Caeca and neck skin samples were received at Microsearch laboratories (Hebden Bridge, West Yorkshire, UK) and cultured onto mCCDA (PO0119A Oxoid Ltd, Basingstoke, UK) and incubated at 42°C in a microaerobic atmosphere for 48 hours. Whole plate sweeps of lowest dilution of *Campylobacter*-positive samples (189 out of 300 pre-vaccination, 679 out of 900 post-vaccination) were stored in 20% glycerol stocks and stored at -80°C. Samples were cultured from frozen glycerol stocks and streaked onto mCCDA (PO0119A Oxoid Ltd, Basingstoke, UK) with CCDA Selective Supplement (SR0155E Oxoid Ltd, Basingstoke, UK) and incubated at 42°C for 48h in a microaerobic atmosphere (85% N_2_, 10% CO_2_, and 5% O_2_) using CampyGen Compact sachets (Thermo Fisher Scientific Oxoid Ltd, Basingstoke UK). Single colonies were isolated and streaked onto mCCDA (PO0119A Oxoid Ltd, Basingstoke, UK) without supplement and incubated for 48h at 42°C in the same microaerobic conditions. A single colony from each plate was then sub-cultured onto Mueller-Hinton (MH) agar (CM0337 Oxoid Ltd, Basingstoke, UK) and grown for an additional 48h at 42°C ready for DNA extraction. Enumeration of cfu/g were calculated for each caeca and neck skin sample and are located in S6 Table.

### Genome sequencing and assembly

DNA was extracted using the QIAamp DNA Mini Kit (QIAGEN, Crawley, UK), according to manufacturer’s instructions. DNA was quantified using a Nanodrop spectrophotometer before sequencing. Genome sequencing was performed on an Illumina MiSeq sequencer using the Nextera XT Library Preparation Kit with standard protocols. Libraries were sequenced using 2 × 300 bp paired end v3 reagent kit (Illumina), following manufacturer’s protocols. Short read paired-end data was assembled using the *de novo* assembly algorithm, SPAdes (version 3.10.0 35)^63^. The average number of contigs was 355.10 (range: 28-1333) (pre-vaccination) and 863.58 (range: 273-2703) (post-vaccination) for an average total assembled sequence size of 1.66 Mbp (range: 1.23 Mbp – 1.98 Mbp) (pre-vaccination) and 1.48 Mbp (range: 1.03 Mbp – 2.74 Mbp) (post-vaccination) (S1 Table).

### Phylogenetics and identification of survival-associated genetic elements

To understand the genetic variation of pre-vaccination isolates, a core gene alignment was constructed using MAFFT with default parameters of minimum nucleotide identity of 70% over >50% of the gene and a BLAST-n word size of 20. This was in comparison to reference NCTC11168 genome (accession number: NC_002163.1) for the 136 pre-vaccination isolate genomes combined with an additional 958 genomes belonging to chicken-associated lineages (S2 Table). A maximum-likelihood (ML) phylogeny was constructed using FastTree version 2.1.8 and the Generalised time-reversibly (*GTR*) model of nucleotide evolution^64^ (Fig 1). A previous genome-wide association study (GWAS) comparing the variation of isolate composition of *Campylobacter* on the farm, meat and in clinical cases identified genetic elements associated with survival through poultry processing^41^. The elements were divided into seven functional categories: ‘transport and binding proteins’, ‘cell envelope’, ‘energy metabolism’, ‘translation’, ‘amino acid biosynthesis’, ‘hypothetical’ and ‘other’ (S4 Table). The 70 x 30 bp genetic elements with the strongest association with survival (p value ≥ 1.00 x 10^-6^) (S4 Table) were used to identify isolates with the potential to “survive” through factory processing in the pre-vaccination sampling dataset (136 genomes). The 30bp “words” were blasted against the pre-vaccination genomes to identify matches at 100% similarity. A presence-absence matrix was returned for each genome (Fig 1B).

### Autogenous vaccine production

The four *C. jejuni* vaccine isolates were recovered from -80°C storage by inoculating Columbia blood agar plates supplemented with yeast extract (CBA+YE) provided by Southern Group Laboratory, Corby UK (SGL) with a Microbank bead (Pro-Lab Diagnostics, Toronto Canada) and incubating at 37°C for 48 hours in a microaerobic atmosphere (85% N_2_, 10% CO_2_, and 5% O_2_) using CampyGen Compact sachets (Thermo Fisher Scientific Oxoid Ltd, Basingstoke UK). For each vaccine isolate a lawn was prepared by streaking 1 cm^2^ of colonies to a lawn on CBA+YE plates and incubating at 37°C for 48 hours under the same microaerobic conditions. Each batch of 120 plates was inoculated by harvesting the lawn plate into 400 ml of Nutrient broth + yeast extract (SGL) using a swab and pipetting 1.5 ml of this cell suspension onto 120 CBA+YE plates. This was done rapidly to minimise the time the cells were exposed to air. The plates were incubated for 48 hours at 37°C inside 26 litre aluminium tins containing CampyGen sachets (6x 3.5 l and 2x 2.5 l sachets) and sealed with duct tape. The cultures were harvested by scraping the cell mass into 190 ml of saline containing 0.475% formaldehyde and 0.02% thiomersal.

One batch of each isolate was grown in fermenters. Specifically, the Electrolab (Tewkesbury UK) Fermac 320 bench-top bioreactor control system with 10 litre vessels (working volume 7 litres). The *C. jejuni* isolates were recovered from -80°C storage by inoculating a CBA+YE plate with a Microbank bead and incubating at 37°C for 48 hours in a plastic pouch containing a CampyGen Compact sachet. For each isolate 4 lawns were prepared by streaking 2-3 colonies to lawns on CBA+YE plates and incubating at 37°C for 24 hours in a plastic pouch containing a CampyGen Compact sachet. Each fermenter was inoculated by harvesting the 4 lawns into 10 ml of Nutrient broth (SGL) using swabs and injecting this cell suspension into the inoculation septa of the fermenter. This was done rapidly to minimise the time the cells were exposed to air. The media, 7,000 ml of nutrient broth + yeast extract (SGL), was pumped into the vessel, pre-warmed and incubated for at least 24 hours prior to inoculation to check for sterility. The vessel was stirred at 300 rpm using six blade Ruston impellers. To prevent excessive foaming 10 ml of Antifoam C (Sigma) solution in water (1:4) was injected into the vessel at the time of inoculation. Each fermenter batch culture was grown for 48 hours. A blend of 41,400 ml of finished vaccine product (FVP) was prepared by combining antigen from both plate and fermenter grown batches to give a final cell count for each isolate of 5x10^8^ cells/ml of FVP. The antigen (752 ml) was blended by stirring with saline (19,606 ml), formalin (38.5 ml 39% w/v), thiomersal (303 ml 2%) and Montanide adjuvant ISA 207 R (20,700 ml; Seppic, Puteaux France) to form a low shear water-in-oil-in-water (W/O/W) emulsion. A batch of 80x 500 ml bottles was produced and shipped to the poultry company for vaccination.

### Vaccination protocol and antibody production

A whole farm (4 houses) of Ross breeder chickens (∼40,000 birds) were hyper-immunised by intramuscular injection with the oil-based autogenous vaccine with two doses of 0.5 ml at 14 and 18 weeks of age (May 2018) (Fig 2B and A). Blood samples were wing bled from 30 vaccinated and 30 control (non-vaccinated) breeder birds after the second dose of vaccine was administered (18 weeks of age). This occurred at three time intervals: (i) immediately after second immunisation (10 samples); (ii) ∼35 weeks of age (10 samples); (iii) ∼48-60 weeks of age (10 samples, peak immunity). The same number of eggs were collected at matched time intervals from vaccinated and control cohorts (Fig 2B). When vaccinated broiler breeder flocks had reached peak vaccine immunity age, blood samples were taken from the wing vein of the broiler progeny of vaccinated and unvaccinated breeder birds to monitor the effects of vaccine IgY. Ten samples were also taken at three time intervals: (i) ∼2 days after hatch; (ii) 2 weeks old; (iii) prior to slaughter (∼37 days old) (Fig 2B).

### Antigen production from whole cells

Acid-soluble surface proteins were extracted from the *C. jejuni* vaccine isolates using established protocols (McCoy et al., 1975). Cells were harvested, washed twice in phosphate-buffered saline (PBS) and re-suspended in 0.2 M glycine-HCl (pH 2.2) (25 ml/g wet weight of cells) for 1 hour at room temperature while rolling. The mixture was centrifuged for 10000g x 20 mins (1 hour at 4000rpm) and the supernatant was dialyzed over night at 4°C using Slide-A-Lyzer^TM^ G2 dialysis cassettes against PBS and stored in aliquots at -20°C. Protein concentrations for all extracts were determined using the Pierce™ BCA Protein Assay Kit.

### Enzyme-linked immunosorbent assay (ELISA)

Serum and egg yolk “anti-survivor” *C. jejuni* IgY antibodies were monitored using Enzyme-linked immunosorbent assays (ELISA). Acid-extracted (glycine hydrochloride) antigens of the *Campylobacter* vaccine isolates were used as the capture protein, serum/egg yolk samples as the target antibody and an anti-chicken IgY as the detection body. Microtiter plates (Polysorb; Nunc, Denmark) were coated with 2 µg/ml of acid-extracted surface proteins, both at 100ul/well of *C. jejuni* isolates 7916660, 7930870, 7939216, 7939276 in coupling buffer (0.05M Na2CO3, pH 9.6) overnight at 4°C. The wells were washed three times with ELISA wash buffer (0.85% [w/v] NaCl, 0.05% [v/v] Tween 20) and then probed with 100µl chicken sera diluted (1:200) in ELISA diluent (ELISA wash containing 1% [w/v] bovine serum albumin and 0.5% [w/v] Tris pH 7.4) for 2 hours at 37°C. A preliminary ELISA (results not shown) testing a range of dilutions of sera from 9 weeks old *C. jejuni* colonised and control birds was performed using a single dilution factor (1:200) chosen as a result of good discrimination between the two groups. The wells were washed as before and then probed with 100µl rabbit anti-chicken IgG conjugated to horseradish peroxidase (HRP) (Sigma Ltd.) diluted 1:3000 in ELISA diluent for 30 min at 37°C. After washing, 100µl of 3,3’,5,5’-tetramethylbenzidene (Cambridge Veterinary Services, Cambridge, UK) was added to each well and the microplate incubated at room temperature. The reaction was stopped after 10 min by the addition of 50 µl 2 M H_2_SO_4_. The absorbance was read at 450nm with wavelength correction at 620nm on a Tecan Spark 10M microplate reader.

### Statistical analysis

Statistical analyses were performed using the rstatix package. Overall difference of means (global test) between pre-vaccine, post-vaccine control and post-vaccine vaccinated groups were carried out using the Kruskal-Wallis test (non-parametric test for not normally distributed data). Pairwise comparisons between means to test for significance (cut-off = *P* ≤ 0.05, all *P*-values shown) were performed using non-parametric Mann-Whitney U-tests.

### Modelling post-vaccination strain replacement

A list of 47 potentially antigenic *Campylobacter* genes was compiled (S5 Table). Genetic variation at these loci was characterised in pre- and post-vaccination *Campylobacter* genomes contextualised with the additional 958 chicken-associated genomes (Fig 1, S2 Table). Mapped nucleotide sequences were translated to amino acids and aligned using Clustal Omega (version 1.2.4)^65,66^. Pairwise hamming distances of amino acid content between genomes were estimated using psdm (version 0.1.0)^67^ with gaps included as differences, and a distance matrix created. Based on the vaccine trial data, a logistic regression model-based approach was used to predict the expected population composition post-vaccination. The post-vaccination survival probabilities of the isolates were estimated by combining the relative frequencies of vaccine inclusion isolates belonging to a pre- vs post-vaccination population (intercept of the model), with the minimum antigenic distances of the pre- and post-vaccine genomes to one of the four vaccine isolates (model covariate). A range of meaningful values of the model parameter theta were assessed for the best fit. An appropriate theta (odds ratio) was chosen and applied to the distances from the large context collection dataset to the four vaccine-inclusion isolates in order to make predictions about the effect of the vaccine antigens against other chicken-associated CCs.

## Author Contributions

JKC, SKS, TW designed the study. JKC, JL and EM performed genomic analysis. MEP performed statistical analysis and modelling. JM carried out the ELISA’s and RML provided guidance on protocols. JKC, BP and MDH sequenced the genomes. PH oversaw vaccination and sampling of poultry. DJH and TSW manufactured the vaccine. JKC and BP drafted the manuscript. JC guided the methodology for the modelling. SKS conceptualised the study.

## Data Availability

Short read genome data are available from the NCBI (National Center for Biotechnology Information) SRA (Sequence Read Archive), associated with BioProject PRJNA1033694 (https://www.ncbi.nlm.nih.gov/bioproject/PRJNA1033694). Assembled genomes and supplementary material are available from FigShare: doi: doi: 10.6084/m9.figshare.24464143. Individual accession numbers and pubMLST IDs can be found in Supplementary Tables 2 and 3.

