## Supplementary figures and images for "Genomic tailoring of autogenous poultry vaccines to reduce *Campylobacter* from farm to fork"

### S1 Fig

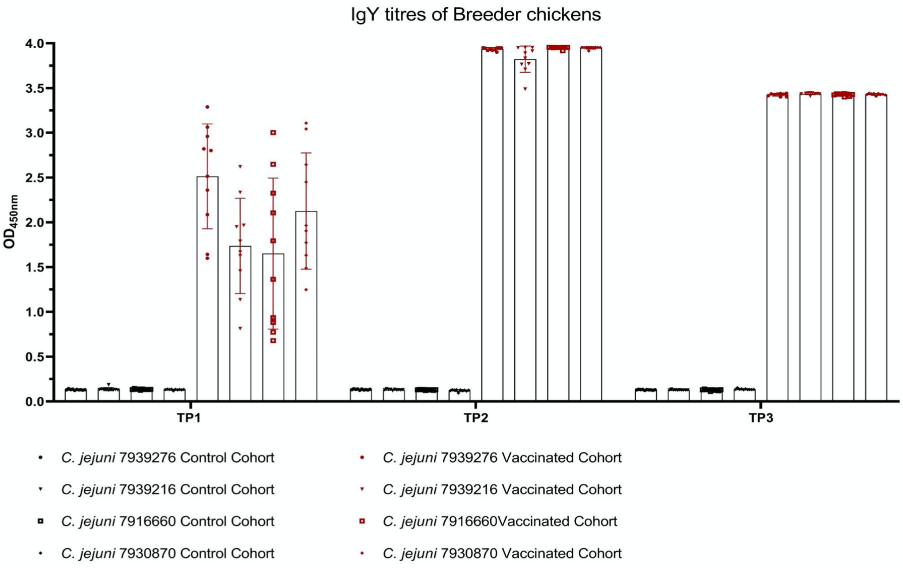

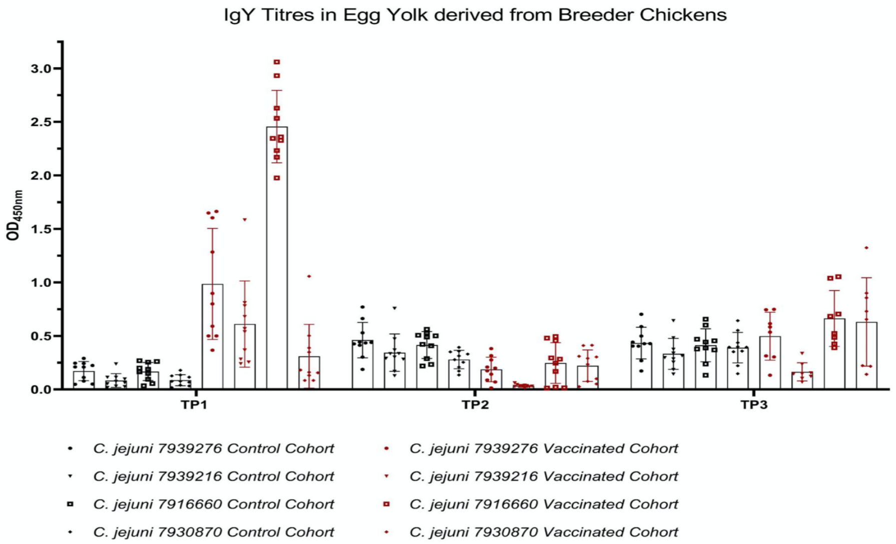

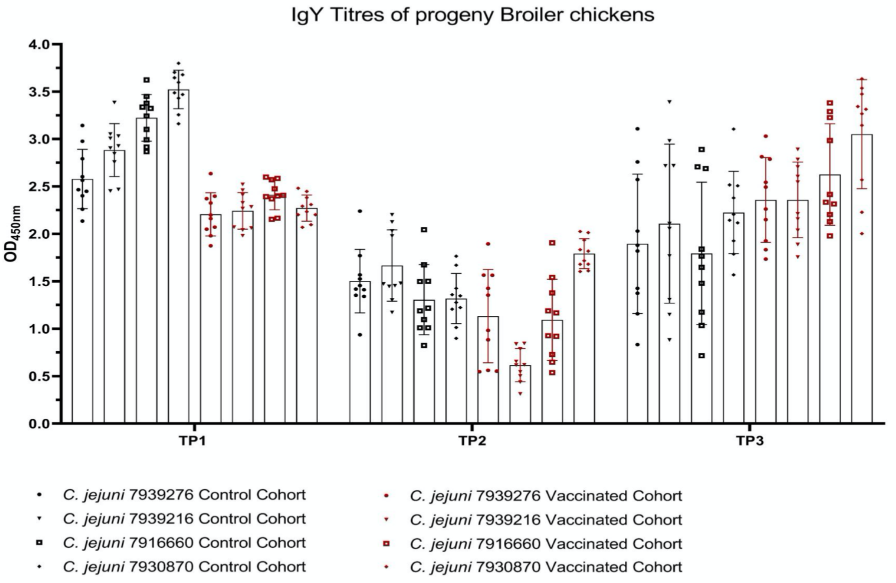

### S4 Fig

A) Odds ratio = 2 (theta = 0.693)

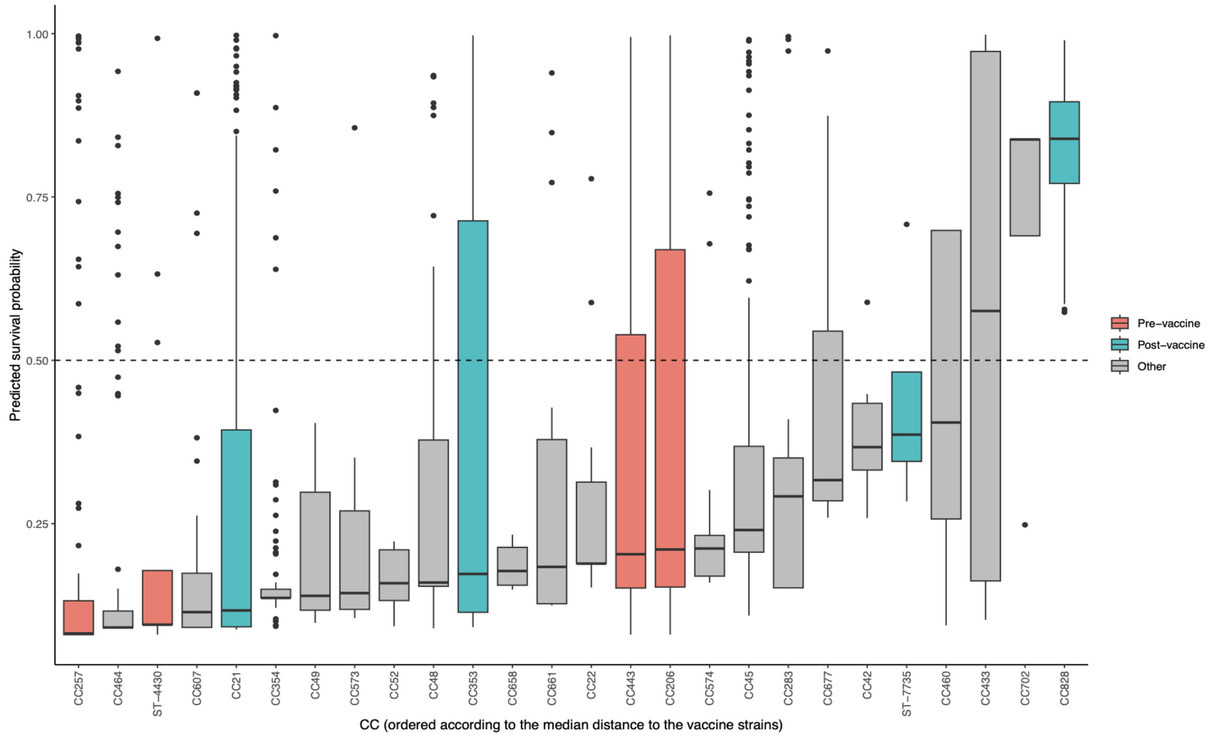


B) Odds ratio = 3 (theta = 1.099)

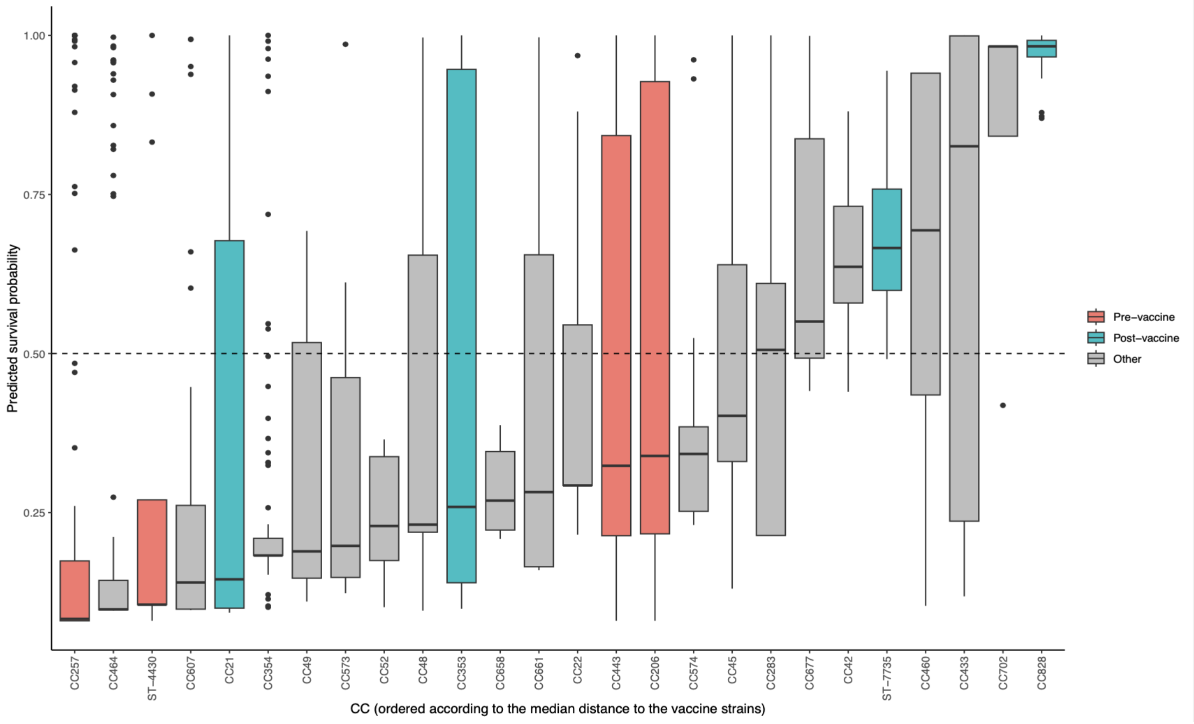

### S5 Fig

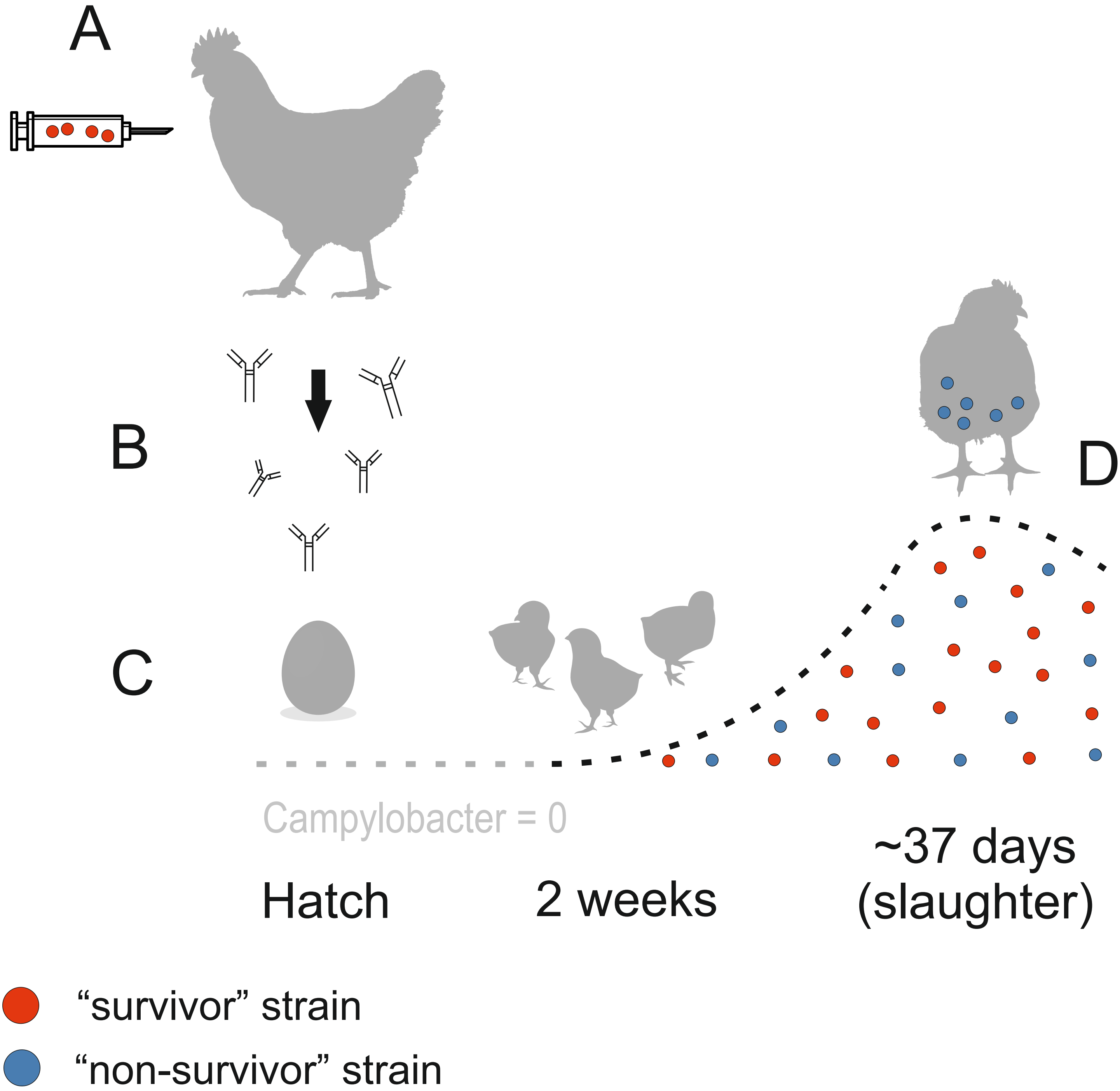
