## Supplementary material for "Genomic tailoring of autogenous poultry vaccines to reduce *Campylobacter* from farm to fork": S2 Fig

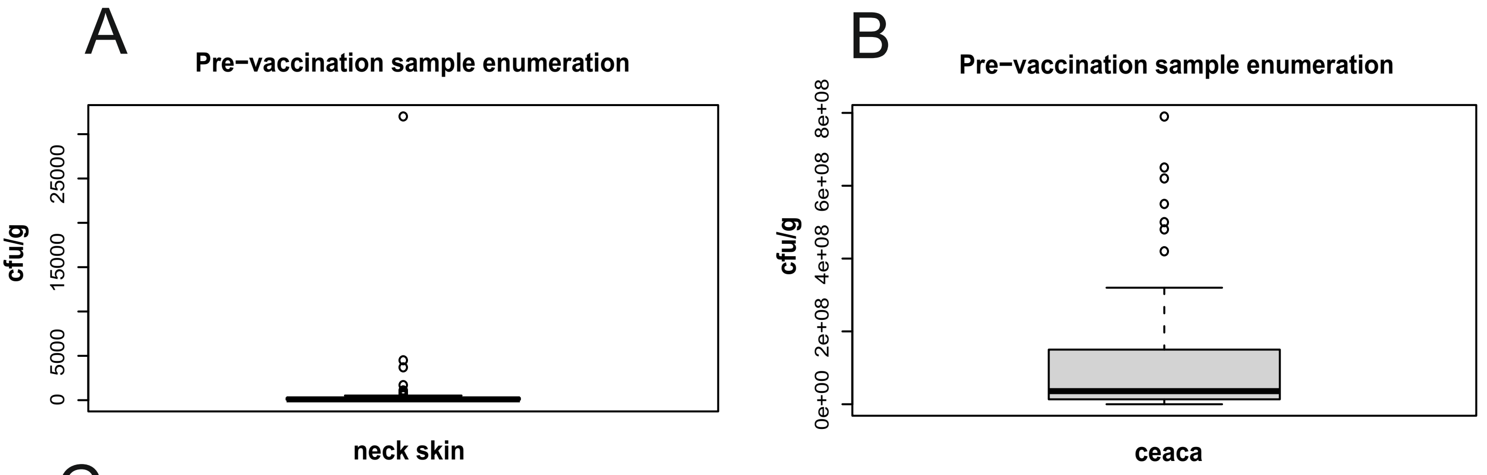


C

| Pre-vac sampling- understanding strain diversity | | | | | | |
|---|---|---|---|---|---|---|
| | Neck skin | | | ceaca | | |
| | Avg. (cfu/g) | No. of samples collected | % Campylobacter +ve | Avg. (cfu/g) | No. of samples collected | % Campylobacter +ve |
| **Farm 1** | 334 | 30 | 83 | 2.05E+08 | 30 | 100 |
| **Farm 2** | 32 | 30 | 27 | 2.95E+06 | 30 | 7 |
| **Farm 3** | 335 | 30 | 70 | 9.54E+07 | 30 | 73 |
| **Farm 4** | 50 | 30 | 63 | 0 | 30 | 0 |
| **Farm 5** | 1389 | 30 | 97 | 2.80E+07 | 30 | 97 |
