## Supplementary material for "Genomic tailoring of autogenous poultry vaccines to reduce *Campylobacter* from farm to fork": S3 Fig

### Post-vaccination sample enumeration

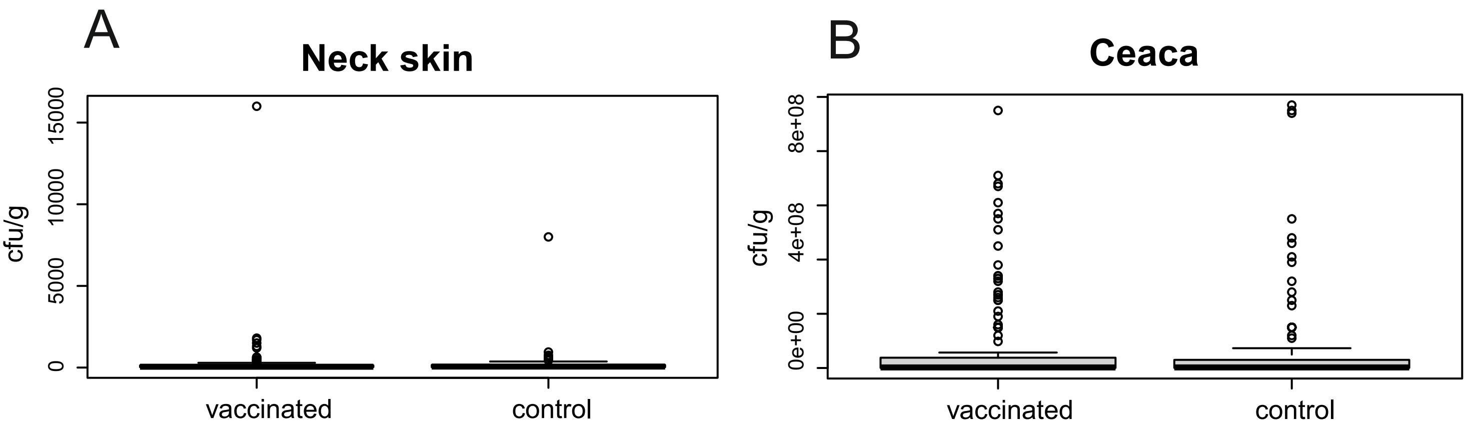


C

| Post-vac sampling - unvaccinated (control) cohort | | | | | | |
|---|---|---|---|---|---|---|
| | Neck skin | | | ceaca | | |
| | **Avg. (cfu/g)** | **No. of samples collected** | **% Campylobacter +ve** | **Avg. (cfu/g)** | **No. of samples collected** | **% Campylobacter +ve** |
| **Farm 1** | 662 | 30 | 53 | 3.18E+06 | 30 | 100 |
| **Farm 2** | 176 | 30 | 83 | 3.07E+07 | 30 | 100 |
| **Farm 3** | 113 | 30 | 40 | 8.18E+05 | 30 | 100 |
| **Farm 4** | 100 | 30 | 87 | 1.69E+08 | 30 | 70 |
| **Farm 5** | 47.5 | 30 | 27 | 1.53E+08 | 30 | 100 |

D

| Post-vac sampling - Vaccinated cohort | | | | | | |
|---|---|---|---|---|---|---|
| | Neck skin | | | ceaca | | |
| | **Avg. (cfu/g)** | **No. of samples collected** | **Campylobacter +ve** | **Avg. (cfu/g)** | **No. of samples collected** | **Campylobacter +ve** |
| **Farm 1** | 77 | 60 | 70 | 9.60E+06 | 60 | 100 |
| **Farm 2** | 3626 | 60 | 98 | 2.40E+07 | 60 | 98 |
| **Farm 3** | 128 | 60 | 45 | 1.17E+06 | 60 | 83 |
| **Farm 4** | 113 | 60 | 33 | 1.83E+08 | 60 | 77 |
| **Farm 5** | 57 | 60 | 50 | 1.19E+08 | 60 | 97 |
